# Restoring Klf9 Expression with Pressure Overload Leads to Metabolic Maladaptation and Early Onset of Heart Failure

**DOI:** 10.64898/2026.07.23.740419

**Authors:** Aishwarya Venkatasubramanian, Chandni Thakkar, Zhi Yang, Ivessa Andreas, Nazish Sayed, Maha Abdellatif, Danish Sayed

**Affiliations:** From the Department of Cell Biology and Molecular Medicine, Rutgers New Jersey Medical School, Newark, New Jersey 07103

## Abstract

Klf9 is a cardiac-enriched transcription factor of the Krüppel-like factor (Klf) family. Klf9 levels decrease during cardiac hypertrophy; however, no studies have examined its transcriptional targets or role in the progression of hypertrophy. Here, we report genome-wide differential Klf9 occupancy during cardiac hypertrophy, with a predominant enrichment at the metabolic gene promoters. Further, using conditional Klf9 knock-in mice subjected to pressure overload for 1 or 2 weeks, we show that restoring Klf9 expression initially inhibits hypertrophy but later leads to early-onset heart failure. We conclude that a decrease in Klf9 is required for metabolic adaptations that support the development of compensatory hypertrophy.

## Introduction

Krüppel-like factors (Klf) have been shown to regulate various cellular processes, including metabolism, differentiation, and proliferation, with implications in development and diseases of the cardiovascular, nervous, respiratory, and immune systems [1, 2]. KLF family members share a conserved C-terminal DNA-binding domain; however, the variable regulatory N-terminal domain, based on its associating factors, can have different effects on target gene expression. Klf9, a member of the Klf family, is enriched in the heart and has been predicted to function as a transcriptional repressor [1] and to exhibit unique genomic targeting compared to other Klfs.

We recently reported the detrimental effects of constitutive expression of Klf9 on postnatal cardiac development, where the αMHC-Cre-Klf9 knock-in (*c*Klf9KI) mice develop spontaneous dilated cardiomyopathy and death by 8-12weeks of age. Characterization of *c*Klf9KI mice revealed dysregulation in metabolic genes as early as 7 days after birth, resulting in a metabolic crisis in the postnatal hearts, which subsequently led to cardiac dysfunction and failure [3]. Others and we have shown that cardiac Klf9 expression decreases during cardiac hypertrophy and failure [3-5], and a similar decrease has been observed in human heart failure [5, 6]. This suggests that changes in Klf9 levels contribute to the stress-induced progression of heart failure.

In our previous study, we were the first to report the direct transcriptional targets of Klf9 in isolated neonatal cardiomyocytes. We show that Klf9 primarily associates with the promoters of metabolic genes and plays a role in metabolic adaptations, as well as serving as one of the feedforward mediators of Glucocorticoid receptor (GR) signaling [7].

Few studies have examined its function in the adult heart. Recently, two studies reported conflicting roles of Klf9 during cardiac hypertrophy: one showed that Klf9 induction protects mitochondrial function and inhibits Angiotensin II-induced hypertrophy [5], and the other showed that Klf9 inhibition rescues Isoproterenol-induced cardiac hypertrophy through its regulation of the long noncoding RNA (lncRNA) UCA1/p27 axis [4]. However, no studies have examined genome-wide Klf9 transcriptional targets in adult hearts under basal conditions or during cardiac hypertrophy, nor their contribution to the progression of heart failure.

## Results

Here, we report the differential Klf9 occupancy on cardiac gene promoters in hearts undergoing sham and transaortic constriction (TAC) operations for 2 weeks (GSE287642), a progressive state characterized by cardiac hypertrophy but no dysfunction at this stage (**Fig. 1A**). Klf9 occupancy data identify 6309 and 5675 genes with 4655 and 4121 Klf9 intervals in Sham and TAC hearts, respectively. Interestingly, 87.09% (4054 of 4655) and 86.9% (3581 of 4121) intervals are within the promoter regions (-7500/+2500bp of NCBI gene start), of which 71.08% (3309 of 4655) and 70.08% (2888 of 4121) intervals are within 500bp of the transcription start site in sham and TAC hearts, respectively (**Fig. 1B** and **1C**). To compare the Klf9 peak metrics between sham and TAC hearts, overlapping and unique intervals are grouped into merged regions. We identify 4645 and 4073 merged regions in Sham and TAC hearts, respectively, of which 3480 were common to both conditions (**Fig. 1D**). For further analysis, we select genes with MaxTags ≥ 50, excluding those with low signal values, as these are expected to be more variable and therefore yield unreliable, often exaggerated ratios. As expected, we observe more merged regions with a decrease (1167 merged regions with a log2FC of ≤ 0.1, ∼10% decrease), versus merged regions with an increase (499 merged regions with a log2FC of ≥0.1, ∼10% increase) in Klf9 occupancy in TAC hearts compared to sham (**Fig. 1E**). The specificity of the Klf9 ChIP-seq is also validated by Homer Motif analysis of the top 2500 peaks (**Fig. 1F**).

**Figure 1.**
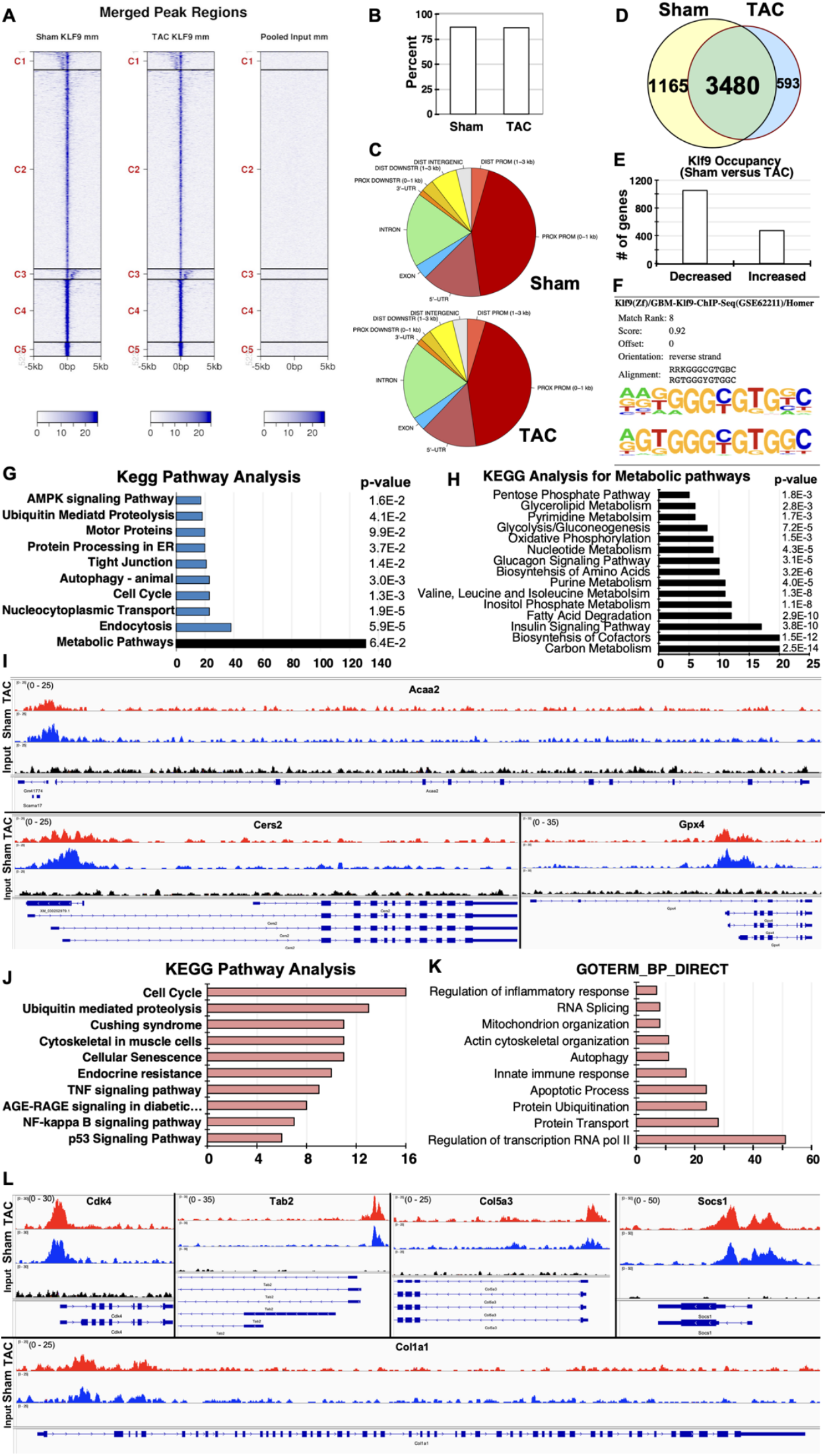
Decrease in Klf9 genomic occupancy at the promoters of metabolic genes during cardiac hypertrophy. **A.** Heatmap showing tag distribution for Klf9 across merged peak regions (values in z-axis/color, active regions in y-axis) in Sham, TAC hearts, and Input control. The data is presented in 5 clusters (default: C1-C5) and sorted. **B**. The graph shows the percentage of Klf9 intervals within the promoter regions of annotated genes. **C**. Pie charts show the location of the Klf9 peaks relative to genomic annotations in the hearts of adult mice subjected to sham or TAC operations. **D**. The Venn diagram shows the number of unique and overlapping Klf9 merged regions in Sham and TAC hearts. **E**. The graph shows the number of genes with differential regulation of Klf9 occupancy in sham versus TAC hearts. **F**. Enriched motif identified using Homer Motif analysis using the 200 bp surrounding the summit of the top 2,500 peaks (based on MACS2 p-values). **G**. Graph lists the number of genes in the functional groups as categorized by KEGG pathway analysis using the DAVID algorithm of genes that show a decrease in Klf9 occupancy in TAC hearts compared to Sham hearts. **H**. The graph lists the number of genes in the sub-metabolic group of the metabolic group as categorized by KEGG pathway analysis. **I**. Plots show Klf9 occupancy on adult (sham) mouse hearts and input samples, as viewed on the integrated genome browser. The X-axis represents chromosomal coordinates, reference sequence, and gene structure. The Y-axis shows the peak value for Klf9 genomic occupancy along the gene structure. **J**. Graph lists the number of genes in the functional groups as categorized by KEGG pathway analysis using the DAVID algorithm of genes that show an increase in Klf9 occupancy in TAC hearts compared to Sham hearts. **K**. The graphs list the number of genes in the functional categories as analyzed by GOTERM using the DAVID algorithm. **L**. Plots show Klf9 occupancy on adult (sham) mouse hearts and input samples, as viewed on the integrated genome browser. The X-axis represents chromosomal coordinates, reference sequence, and gene structure. The Y-axis shows the peak value for Klf9 genomic occupancy along the gene structure.

These data confirm that Klf9 associates with cardiac gene promoters and show a decrease in Klf9 chromatin binding in TAC-operated hearts compared to Sham-operated hearts. KEGG pathway analysis of genes showing decreased Klf9 promoter occupancy lists metabolic pathways at the top, along with endocytosis, nucleocytoplasmic transport, autophagy, tight junctions, motor proteins, ubiquitin-mediated proteolysis, and AMPK signaling (**Fig. 1G**). Further analysis of metabolic pathway-related genes highlighted roles in Carbon metabolism, Biosynthesis of cofactors, Insulin signaling pathways, Fatty acid metabolism, Biosynthesis of Amino Acids, Nucleotide metabolism, Glucagon signaling, Oxidative phosphorylation, Glycolysis/gluconeogenesis, Glycerolipid metabolism, and the Pentose Phosphate pathway (**Fig. 1H**). Fig. 1I shows Klf9 occupancy data for selected representative genes, as viewed in the integrated genome viewer (**Fig. 1I**). These data from sham and TAC hearts validate our previous findings in isolated neonatal cardiomyocytes [7] and Klf9KI hearts [3], where central downstream effects on metabolic pathways for cellular homeostasis are observed with increased Klf9 expression levels, either through GR activity or constitutive postnatal expression. On the other hand, genes that show increased Klf9 occupancy are categorized as involved in the cell cycle, protein degradation, Cushing syndrome, cellular senescence, endocrine resistance, and p53 signaling pathways (**Fig. 1J**). GOTERM analysis of these genes also identifies those involved in inflammation, mitochondrial organization, and apoptosis (**Fig. 1K**) and may reflect contributions from non-cardiomyocytes, such as proliferating fibroblasts and infiltrating immune cells. Fig. 1L shows Klf9 occupancy data for selected representative genes, as viewed in the integrated genome viewer (**Fig. 1L**). These results confirm that Klf9 associates with the promoters of metabolic genes in basal hearts and regulates metabolic adaptations in response to physiological variations, thereby maintaining cellular homeostasis.

Based on our previous publications of GR-Klf9 axis in neonatal cardiomyocytes [7], *c*Klf9KI in postnatal hearts [3] and the Klf9 occupancy data from Sham and TAC-operated hearts, led us to two questions: 1) Is a decrease in Klf9 expression required for the metabolic adaptations in the heart to compensate for increasing work overload, and development of compensatory hypertrophy or 2) does a decrease in Klf9 results in metabolic maladaptation that contributes to the progression of pathological cardiac hypertrophy, and restoring the levels could restrict or delay the development of heart failure.

Therefore, to examine the impact of the decrease in Klf9 expression during cardiac hypertrophy, we subjected conditionally (tamoxifen) induced Klf9KI (Klf9KI) mice to Sham or TAC operations for two time points (1 week and 2 weeks) (**Fig. 2A**). As expected, we observe a 3- to 5-fold increase in Klf9 transcript levels in Klf9KI hearts subjected to Sham or TAC hearts compared to their respective controls (**Fig 2B**), which normalized the protein expression of Klf9 to endogenous sham levels in TAC-operated hearts (**Fig. 2C**). Interestingly, the physical and functional characterization of these hearts reveals an initial inhibition of compensatory hypertrophy at 7 days post-TAC in Klf9KI mice; however, at 14 days post-TAC, they show an earlier onset of decompensation associated with cardiac dysfunction than MCM control mice. Inhibition of compensatory hypertrophy after 7 days post-TAC is evident from normalized heart weight to body weight or tibia length (**Fig 2D**), a decrease in interventricular septum (IVS) (**Fig. 2E**), and left ventricular posterior wall thickness (LVPW) (**Fig. 2F**), during systole and diastole. These changes are accompanied by decreases in ejection fraction (EF) and fractional shortening (FS) in Klf9KI+TAC compared with MCM+TAC hearts; however, no change in left ventricular internal dimensions is observed at 7 days post-TAC. At 2 weeks, along with a decrease in IVS, LVPW, percent EF, and FS (**Fig. 2H**), we observe a significant increase in left ventricular internal dimensions (LVID) during systole and diastole (**Fig. 2G**), suggesting an early onset of cardiac dysfunction and dilatation in Klf9KI+TAC (2wks) hearts compared to MCM-TAC. Wheat germ agglutinin (WGA) staining shows a limited increase in individual cardiomyocyte area after 7 days post-TAC in Klf9KI+TAC hearts compared with MCM+TAC, suggesting that TAC-induced cardiomyocyte hypertrophy is inhibited. However, by day 14, cardiomyocytes in Klf9KI+TAC hearts show increased area compared with those in MCM+TAC hearts (**Fig. 2I** and **2J**), suggesting exaggerated cardiomyocyte hypertrophy and dilatation. Further, we observe an increase in the ventricular cavity on H&E stain (**Fig. 2K**), interstitial fibrosis (**Fig. 2L** and **2M**), and TUNEL-positive nuclei (**Fig. 2N**), suggesting an increase in cardiomyocyte apoptosis in the Klf9KI+TAC compared to MCM+TAC hearts after 2 weeks. Next, to examine the effects of Klf9 restoration on mitochondrial function, we measure the activities of complexes I, II, and IV in isolated mitochondria from these hearts subjected to 2 weeks of TAC. We observe a significant decrease in Complex I, II, and IV activity in Klf9KI-TAC hearts compared with control MCM-TAC hearts (**Fig 2O** and **2P**), suggesting a reduction in mitochondrial function. These data indicate that restoring Klf9 expression levels during pressure overload-induced compensatory cardiac hypertrophy promotes the early onset of cardiac dysfunction and the progression of pathological cardiac hypertrophy compared to MCM-hearts. These results further suggest that a decrease in Klf9 expression, coinciding with compensatory hypertrophy and the absence of cardiac dysfunction, may be required for the cardiomyocyte metabolic adaptations to pressure overload.

**Figure 2.**
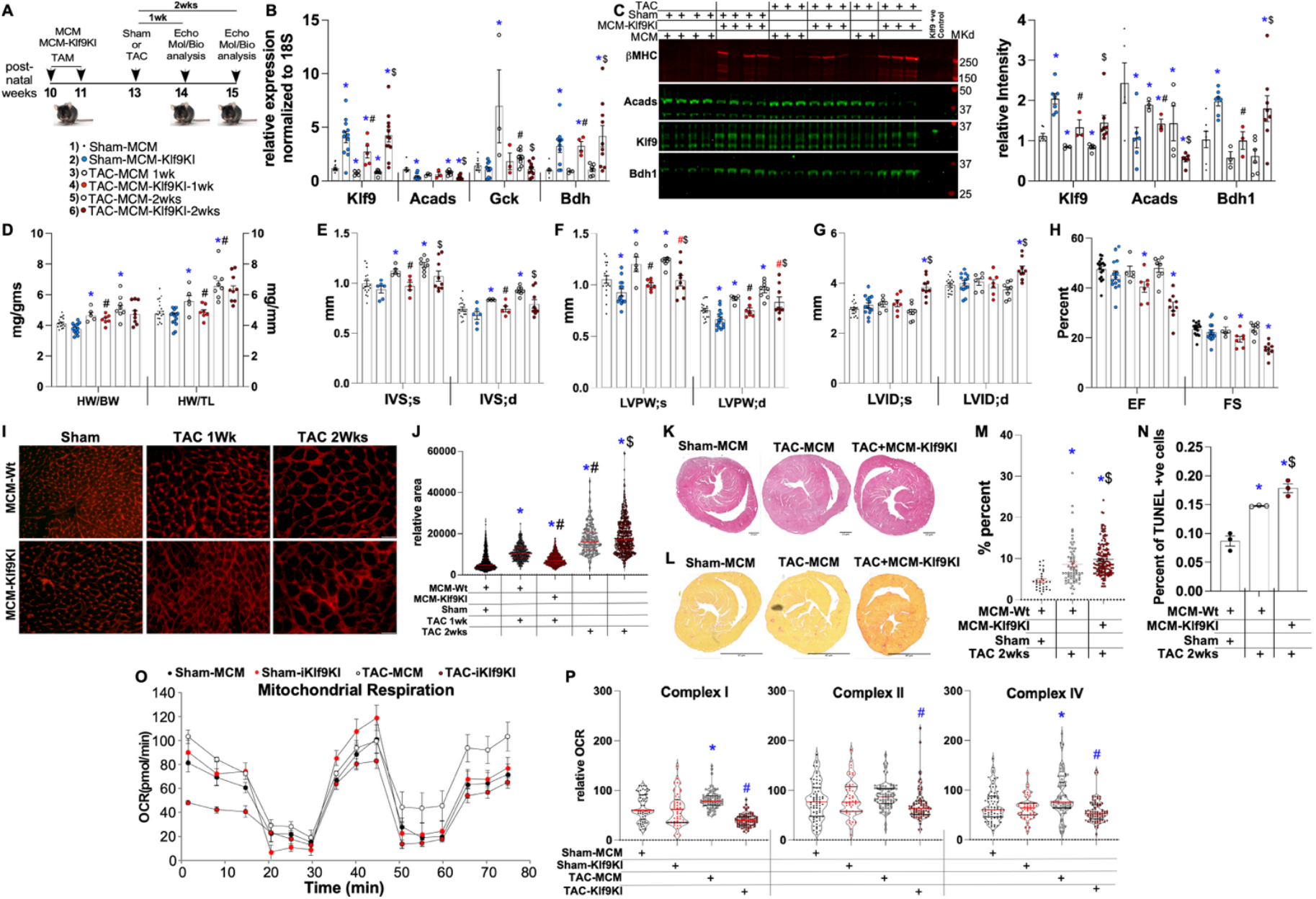
Restoring Klf9 levels during TAC-induced cardiac hypertrophy results in an early onset of cardiac dysfunction. **A.** The schematic shows the experimental design, including details on age groups, genotypes, Tamoxifen administration date, TAC dates, and duration. Below the schematic, the color key with the corresponding samples is shown for all graphs, in the sequence: Sham-MCM, Klf9KI-MCM, TAC (1wk)-MCM, TAC (1wk)-Klf9KI, TAC (2wks)-MCM, and TAC (2wks)-Klf9KI. **B**. The graph shows the relative expression of the selected genes, normalized to the internal control 18S. **C**. Western blot showing the expression levels of indicated proteins, with the quantified graph normalized to the internal control. **D** - **H**. The graphs represents heart weight (HW) normalized to body weight (BW) or tibia length (TL), interventricular septum (IVS), left ventricular posterior wall (LVPW) thickness, left ventricular internal dimensions (LVID) during systole (s) and diastole (d) and percent ejection fraction (EF) and fractional shortening (FS) in Mer-Cre-Mer (MCM)-sham, MCM-Klf9KI-sham, MCM-TAC and MCM-Klf9KI-TAC after 1wk and 2 weeks of sham or TAC operations. **I** and **J**. Wheat germ agglutinin (WGA) staining showing individual cardiomyocyte area in the MCM-Wt and MCM-Klf9KI hearts after Sham or TAC operations for 1wk or 2 weeks, as indicated. The graph represents the relative quantified area from at least 3 independent hearts for each group. **K** and **L**. Heart sections from MCM-Sham or MCM-Klf9KI subjected to Sham or TAC (2 weeks) were stained with H&E and PASR. **M**. The graph represents the percent fibrosis in MCM-Sham, MCM-TAC (2wks) and MCM-Klf9KI-TAC (2 wks) hearts, as indicated. **N**. The graph shows the percentage of TUNEL-positive nuclei in heart sections from the indicated hearts. **O** and **P**. Mitochondria were freshly isolated from the hearts of Sham, Sham-MCM-Klf9KI, TAC-MCM-2Wks and TAC-MCM-Klf9KI-2wks mice and plated in mitochondrial assay buffer containing pyruvate, malate, and FCCP. The OCR (pmol/min, y-axis) over time (min, x-axis) was measured using the Seahorse XFe96 analyzer before and after the sequential injections of rotenone (Rtn), succinate (succ), antimycin A (AA), and Ascorbic acid + TMPD (Asc/TMPD), at 2-time points for each injection, as seen on the curve. Violin plots on the right show the relative OCRs for each injection, graphed separately for Basal (Complex I), Complex II, and Complex IV-mediated respiration. The data were normalized to mitochondrial protein per well. For all graphs, error bars represent SEM, * is p<0.05 compared to respective Wt-Cre, # is p<0.05 compared to TAC (1wk), and$ is p<0.05 compared to TAC (2 wk) (calculated using unpaired, 2-tailed Student’s t-test), n=3-8 independent hearts, as indicated.

## Discussion

Klf9, a member of the zinc finger transcription factor family, and its expression is decreased during cardiac hypertrophy and heart failure. While the roles of other Klf family members, such as Klf15 and Klf4, have been investigated in cardiovascular disorders, the functional role of Klf9 in cardiac hypertrophy and failure remains controversial and poorly understood.

In this study, we demonstrate the previously unexplored functional significance of Klf9 in pressure overload-induced cardiac hypertrophy. Our Klf9 ChIP-seq data show decreased Klf9 occupancy at the promoters of metabolic genes in TAC-operated hearts compared with sham. Notably, genes from pathways involved in glycolysis and fatty acid metabolism exhibited reduced Klf9 occupancy to their promoters. We show the differential regulation of two representative genes from these pathways: Acads (or SCAD), a short-chain acyl-CoA dehydrogenase, and Glucokinase (Gck), which are decreased in Klf9KI-TAC hearts compared to MCM-TAC hearts. Although glucokinase (Gck) is enriched in the liver, and hexokinase catalyzes the first critical step in glycolysis [8], a recent study in Circulation shows that glucokinase is expressed in the heart. An increase in Gck transcription reduces apoptosis and improves mitochondrial respiration and glycolysis in cardiomyocytes during ischemia-reperfusion injury [9]. As shown in **Fig 2B**, exogenous Klf9 expression with TAC reduces Acads, thereby hampering fatty acid oxidation, and also inhibits TAC-induced upregulation of Gck and dependent glycolysis. This suggests that Klf9-dependent transcription could regulate the cardiometabolic remodeling and energy homeostasis associated with cardiac hypertrophy.

Our data shows that restoration of Klf9 expression, while initially restricting cardiac hypertrophy, led to an accelerated ventricular dilatation, systolic dysfunction, and cardiac dysfunction by 14 weeks, suggesting an early onset of heart failure. A recent study on hypertrophic obstructive cardiomyopathy also showed that Klf9 downregulation attenuated hypertrophic growth by regulating the UCA1/p27 axis [4]. Consistent with our findings, Klf9 overexpression has been implicated in exacerbating cardiac hypertrophy in diabetic cardiomyopathy by inhibiting Nrf2 signaling [10]. Importantly, the cardiac phenotype observed in our TAC-operated Klf9KI mice was accompanied by impaired mitochondrial function, as indicated by reduced mitochondrial complex activity. This is particularly intriguing, as it has been reported that Klf9 regulates transcription of the PPARγ pathway [10, 11], which can affect downstream pathways involved in maintaining energy homeostasis, such as glucose and lipid metabolism [12], aligning with our transcriptomics and functional data. Previous non-cardiac studies have shown that Klf9 regulates hepatic glucose metabolism [13], represses inflammation, and promotes cytoskeletal remodeling in neurons [14]. However, our findings differ from a recent study that reported cardioprotective effects of Klf9 overexpression against angiotensin II-induced hypertrophy. Additionally, the Klf9-mediated cardioprotective effects in this angiotensin II study pointed towards mitophagy and mitochondrial quality control pathways [5], whereas our study with the pressure-overload model revealed changes in metabolic gene networks. These divergent findings could be explained by differences in the pathological stimuli: Angiotensin II, a neurohormonal stressor, is reported to directly cause mitochondrial injury and mitophagy. Whereas the pressure overload model causes hemodynamic stress, which involves extensive transcriptional and metabolic reprogramming, critical for maintaining the myocardial function [15-17]. Our findings suggest that sustained Klf9 expression during pathological conditions is maladaptive and interferes with the adaptive transcriptional response to TAC-induced cardiac hypertrophy. Furthermore, our data indicate that non-dynamically regulated Klf9 expression disrupts the downstream transcriptional network governing metabolic adaptation and mitochondrial function during pressure overload, contributing to the progression of cardiac dysfunction. We intend to further explore the individual downstream metabolic pathways affected by Klf9 dysregulation in these hearts and to determine the extent to which they contribute to metabolic adaptations to cardiac stress. It will also be interesting to characterize the upstream regulators of the dynamic changes of Klf9 expression during physiological and pathological hypertrophy. We have shown that the GR-Klf9 axis mediates metabolic adaptations in neonatal cardiomyocytes [7]; however, whether GR also regulates Klf9 expression in adult hearts and under stress remains to be examined.

In conclusion, we report that a downregulation of Klf9 is required for metabolic adaptations during compensatory cardiac hypertrophy. Restoring Klf9 levels in mouse hearts undergoing TAC-induced cardiac hypertrophy initially inhibits hypertrophy; however, under persistent pressure overload, it leads to early-onset cardiac dysfunction and heart failure.

## Author contributions

AV performed experiments; CT assisted in the animal colony and mitochondrial experiments; ZY performed the sham/TAC surgeries and Echo on the mice; AI performed the histology and staining; NS assisted in the experimental design; MA assisted in the Seahorse experiments; DS designed experiments, performed data analysis with figures, and wrote the manuscript.

## Acknowledgements

We thank Dr. Sadoshima’s lab, Chair of the Department of Cell Biology and Molecular Medicine, Rutgers New Jersey Medical School, for his support.

## Bibliography

[1] B.B. McConnell, V.W. Yang, Mammalian Krüppel-like factors in health and diseases, Physiol Rev 90(4) (2010) 1337–81.

[2] M.G. Santoyo-Suarez, J.D. Mares-Montemayor, G.R. Padilla-Rivas, J.L. Delgado-Gallegos Quiroz-Reyes, J.A. Roacho-Perez, D.F. Benitez-Chao, L. Garza-Ocañas, G. Arevalo-Martinez, E.N. Garza-Treviño, J.F. Islas, The Involvement of Krüppel-like Factors in Cardiovascular Diseases, Life (Basel) 13(2) (2023).

[3] C. Thakkar, S. Alikunju, A. Venkatasubramanian, D. D’Mello, H. Abbas, Z. Yang, I. Andreas, N. Sayed, M. Abdellatif, D. Sayed, Constitutive expression of cardiomyocyte Klf9 precipitates metabolic dysfunction and spontaneous cardiomyopathy, Cell Signal 136 (2025) 112146.

[4] D. Ding, G. Zhao, KLF9 aggravates the cardiomyocyte hypertrophy in hypertrophic obstructive cardiomyopathy through the lncRNA UCA1/p27 axis, Int J Exp Pathol 106(2) (2025) e12526.

[5] L. Zhang, M. Zhang, J. Huang, J. Huang, Y. Zhang, Y. Zhang, H. Chen, C. Wang, X. Xi, H. Fan, J. Wang, D. Jiang, J. Tian, J. Zhang, Y. Chang, Klf9 is essential for cardiac mitochondrial homeostasis, Nat Cardiovasc Res 3(11) (2024) 1318–1336.

[6] V.S. Hahn, H. Knutsdottir, X. Luo, K. Bedi, K.B. Margulies, S.M. Haldar, M. Stolina, J. Yin, A.Y. Khakoo, J. Vaishnav, J.S. Bader, D.A. Kass, K. Sharma, Myocardial Gene Expression Signatures in Human Heart Failure With Preserved Ejection Fraction, Circulation 143(2) (2021) 120–134.

[7] C. Thakkar, S. Alikunju, N. Niranjan, W. Rizvi, A. Abbas, M. Abdellatif, D. Sayed, Klf9 plays a critical role in GR -dependent metabolic adaptations in cardiomyocytes, Cell Signal 111 (2023) 110886.

[8] K. Wang, M. Shi, C. Huang, B. Fan, A.O.Y. Luk, A.P.S. Kong, R.C.W. Ma, J.C.N. Chan, E. Chow, Evaluating the impact of glucokinase activation on risk of cardiovascular disease: a Mendelian randomisation analysis, Cardiovasc Diabetol 21(1) (2022) 192.

[9] Y. Bei, Y. Zhu, J. Zhou, S. Ai, J. Yao, M. Yin, M. Hu, W. Qi, M. Spanos, L. Li, M. Wei, Z. Huang, J. Gao, C. Liu, P.H. van der Kraak, G. Li, Z. Lei, J.P.G. Sluijter, J. Xiao, Inhibition of Hmbox1 Promotes Cardiomyocyte Survival and Glucose Metabolism Through Gck Activation in Ischemia/Reperfusion Injury, Circulation 150(11) (2024) 848–866.

[10] F. Li, J. Peng, H. Feng, Y. Yang, J. Gao, C. Liu, J. Xu, Y. Zhao, S. Pan, Y. Wang, L. Xu, W. Qian, J. Zong, KLF9 Aggravates Streptozotocin-Induced Diabetic Cardiomyopathy by Inhibiting PPARγ/NRF2 Signalling, Cells 11(21) (2022).

[11] Q. Yan, B. He, G. Hao, Z. Liu, J. Tang, Q. Fu, C.X. Jiang, KLF9 aggravates ischemic injury in cardiomyocytes through augmenting oxidative stress, Life Sci 233 (2019) 116641.

[12] L. Gao, Y. Liu, S. Guo, L. Xiao, L. Wu, Z. Wang, C. Liang, R. Yao, Y. Zhang, LAZ3 protects cardiac remodeling in diabetic cardiomyopathy via regulating miR-21/PPARa signaling, Biochim Biophys Acta Mol Basis Dis 1864(10) (2018) 3322–3338.

[13] A. Cui, H. Fan, Y. Zhang, Y. Zhang, D. Niu, S. Liu, Q. Liu, W. Ma, Z. Shen, L. Shen, Y. Liu, H. Zhang, Y. Xue, Y. Cui, Q. Wang, X. Xiao, F. Fang, J. Yang, Q. Cui, Y. Chang, Dexamethasone-induced Krüppel-like factor 9 expression promotes hepatic gluconeogenesis and hyperglycemia, J Clin Invest 129(6) (2019) 2266–2278.

[14] J.R. Knoedler, A. Subramani, R.J. Denver, The Krüppel-like factor 9 cistrome in mouse hippocampal neurons reveals predominant transcriptional repression via proximal promoter binding, BMC Genomics 18(1) (2017) 299.

[15] A.A. Manolis, T.A. Manolis, A.S. Manolis, Neurohumoral Activation in Heart Failure, Int J Mol Sci 24(20) (2023).

[16] T. Kadoguchi, S. Kinugawa, S. Takada, A. Fukushima, T. Furihata, T. Homma, Y. Masaki, W. Mizushima, M. Nishikawa, M. Takahashi, T. Yokota, S. Matsushima, K. Okita, H. Tsutsui, Angiotensin II can directly induce mitochondrial dysfunction, decrease oxidative fibre number and induce atrophy in mouse hindlimb skeletal muscle, Exp Physiol 100(3) (2015) 312–22.

[17] H. Yang, X. Wang, A Comparative Discussion on the Selection of Cardiac Hypertrophy Models: TAC Surgery Vs Ang II Infusion [Letter], Drug Des Devel Ther 18 (2024) 4563–4564.

